# THOC1 deficiency leads to late-onset nonsyndromic hearing loss through p53-mediated hair cell apoptosis

**DOI:** 10.1101/719823

**Authors:** Luping Zhang, Yu Gao, Ru Zhang, Feifei Sun, Cheng Cheng, Fuping Qian, Xuchu Duan, Guanyun Wei, Xiuhong Pang, Penghui Chen, Renjie Chai, Tao Yang, Hao Wu, Dong Liu

## Abstract

Apoptosis of cochlear hair cells is a key step towards age-related hearing loss. Although numerous genes have been implicated in the genetic causes of late-onset, progressive hearing loss, few show direct links to the proapoptotic process. By genome-wide linkage analysis and whole exome sequencing, we identified a heterozygous p.L183V variant in *THOC1* as the probable cause of the late-onset, progressive, non-syndromic hearing loss in a large dominant family. Thoc1, a member of the conserved multisubunit THO/TREX ribonucleoprotein complex, is highly expressed in mouse and zebrafish hair cells. The *Thoc1* mutant zebrafish generated by gRNA-Cas9 system lacks the C-startle response, indicative of the hearing dysfunction. Both *Thoc1* mutant and knockdown zebrafish have greatly reduced hair cell numbers, while the latter can be rescued by embryonic microinjection of human wild-type *THOC1* mRNA but to significantly lesser degree by the p.L183V mutant mRNA. The *Thoc1* deficiency resulted in marked apoptosis in zebrafish hair cells. Consistently, transcriptome sequencing of the mutants showed significantly increased gene expression in the p53-associated signaling pathway. Depletion of p53 or applying the p53 inhibitor Pifithrin-α significantly rescued the hair cell loss in the *Thoc1* knockdown zebrafish. Our results suggested that *THOC1* deficiency lead to late-onset, progressive hearing loss through p53-mediated hair cell apoptosis. This is to our knowledge the first human disease associated with *THOC1* mutations and may shed light on the molecular mechanism underlying the age-related hearing loss.

**Significance Statement:** For the first time, we found that THOC1 deficiency leads to late-onset nonsyndromic hearing loss. Furthermore, we revealed the hypomorphic THOC1 induced p53-mediated hair cell apoptosis. This is to our knowledge the first human disease associated with THOC1 mutations and may shed light on the molecular mechanism underlying the age-related hearing loss.

## Introduction

Age-related hearing loss affects over 40% of the population older than 65 years[1]. Apoptosis of the inner ear sensory hair cells, which are nonregenerative in mammals, is a key step towards this process[2, 3]. Over the past two decades, more than 40 genes have been implicated in the genetic causes of late-onset, progressive hearing loss (Hereditary Hearing Loss Homepage, http://hereditaryhearingloss.org/). Dysfunction of those genes have been shown to hinder the repair or stability of critical auditory components such as cytoskeleton structure, intercellular junction, fluid homeostasis and synaptic transmission[4]. While such findings provided invaluable resources to gain insights into the molecular basis of the age-related hearing loss, the mechanism underlying the direct cause of the progressive hair cell loss remains elusive. To date, only *TJP2* and *DIABLO*, two genes associated with dominant, late-onset, progressive hearing loss DFNA51 and DFNA64, respectively, have been reported to participate in the apoptosis pathway[5, 6]. Overexpression of the tight junction protein TJP2 due to a genomic duplication was shown to result in decreased phosphorylation of GSK-3β and altered expression of the apoptosis-related genes, increasing the susceptibility of inner ear cells to apoptosis[6]. A missense p.Ser126Leu mutation in the proapoptotic gene *DIABLO* retained its proapoptotic function but renders the mitochondria susceptible to calcium-induced loss of the membrane potential[5].

THOC1 is an essential member of the conserved THO/TREX ribonucleoprotein (RNP) complex that functions in cotranscriptional recruitment of mRNA export proteins to the nascent transcript[7]. Along with the assembly of other THO/TREX proteins, it has been shown to regulate the coordinated gene expression required for cancer development as well as self-renewal and differentiation of embryonic stem cells[8]. Mice homozygous for the *Thoc1* null allele is embryonic lethal, while those with the *Thoc1* hypomorphic allele exhibited a dwarf phenotype with severely compromised gametogenesis[7, 9, 10]. So far, no *THOC1* mutations have been reported in humans and the function of THOC1 in inner ear has not been explored. In this study, we identified a missense mutation in *THOC1* as the probable cause of late-onset, progressive, non-syndromic hearing loss in a large, dominant family. Deficiency of Thoc1 was shown to lead to hair cell apoptosis through the p53-mediated pathway.

## Results

### Clinical characteristics of Family SH

Family SH had at least 15 members affected by adulthood-onset hearing loss within 4 generations (Figure 1a). The nonsyndromic, bilateral hearing loss was most predominant in the high frequencies, beginning mildly during the third decade and gradually progressed to severe-to-profound in the fifth and sixth decades (Supplementary Figure S1). All affected subjects reported tinnitus, while vestibular dysfunction or other clinical abnormalities were not observed. No inner ear malformation was detected by temporal bone computed-tomography (CT) scanning (data not shown).

**Figure 1.**
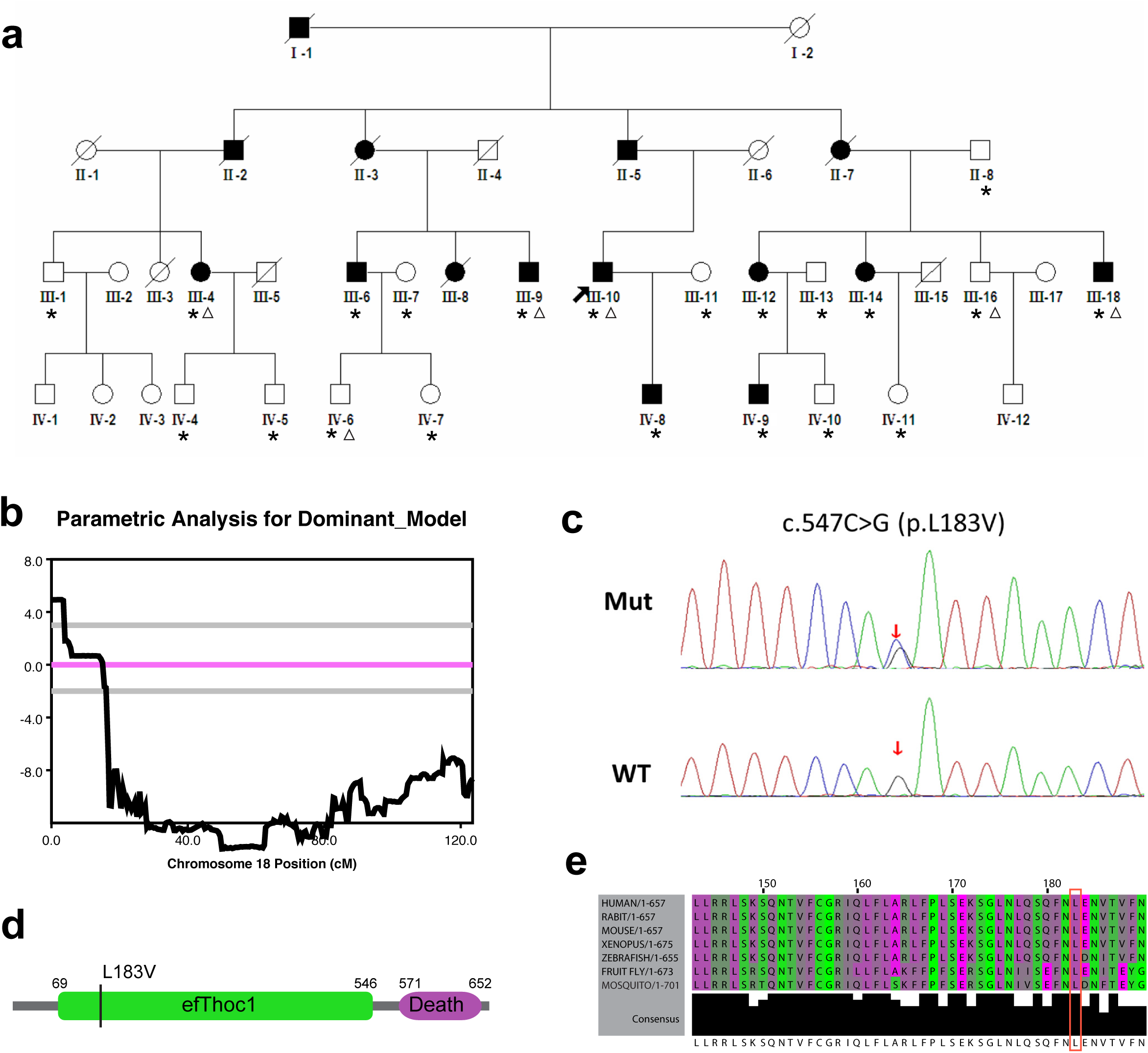
Pedigrees and genotypes of families SH. (a) Pedigrees of families SH. The individuals selected for linkage analysis and whole-exome sequencing was marked with asterisks and triangles, respectively. (b) Logarithm of the odds (LOD) scores of genome-wide linkage analysis for chromosome 18. A maximum LOD score of 4.93 was obtained for marker rs928980. (c) Chromatograms of wild type (WT) and mutant (Mut) sequence for c.C547G (p.L183V). (d) Diagram showing domains of human THOC1 protein and the location of the p.L183V mutation. (e) Multiple sequence alignment of THOC1 showing conservation of the leucine 183 residue.

### Identification of the p.L183V mutation in *THOC1*

Because the onset of the hearing loss was during the third decade for Family SH, only unaffected family members over 40 years old were included in the linkage analysis in this study. Multipoint genome-wide linkage analysis of the 9 affected and 12 unaffected family members (marked with asterisks in Figure 1a) detected a 1.4-Mb critical interval on Chromosome 18p11.3 between markers rs4797697 and rs11080868 (Figure 1b and Supplementary data 1). Maximum logarithm of the odds (LOD) score of 4.93 was obtained for markers rs928980, rs1022177 and rs12709528 (Supplementary Table S1).

Whole-exome sequencing of four affected (III-4, III-9, III-10 and III-18) and two unaffected (III-16 and IV-6) family members (marked with triangles in Figure 1a) identified a total of three candidate variants (Supplementary Table S2). Among them, only the c.C547G (p. L183V) variant in *THOC1* (NM_005131) segregated with the hearing loss in the rest of the family members (Figure 1c). Consistent with the linkage analysis results, the p. L183V variant in *THOC1* is within the 1.4-Mb critical interval. Leu183 resides in the elongation factor (efThoc1) domain of THOC1 and is evolutionarily conserved from human to fruit fly (Figure 1d, e; Supplementary Figure S2). The p.L183V variant was predicted to be pathogenic by computational programs Mutation Taster, PROVEAN and SIFT and was not seen in 1000 Chinese Han normal hearing controls.

### The expression of *thoc1* in mouse and zebrafish hair cells

To further elucidate the role of THOC1 in hearing, we investigated the expression of THOC1 in mouse and zebrafish. Cross-section and whole mount immunostaining of P0 mouse inner ear showed that THOC1 was specifically expressed in inner and outer hair cells (Figure 2a, b) but not in saccule, utricles, spiral ganglion cells and stria vascularis (Supplementary Figure S3). Consistently, whole mount *in situ* hybridization in zebrafish showed that *thoc1* was specifically enriched in the zebrafish developing neuromast, particularly in the hair cells (Figure 2c-e’). These results suggest that Thoc1 conservatively contribute to the formation of hair cells in vertebrates.

**Figure 2.**
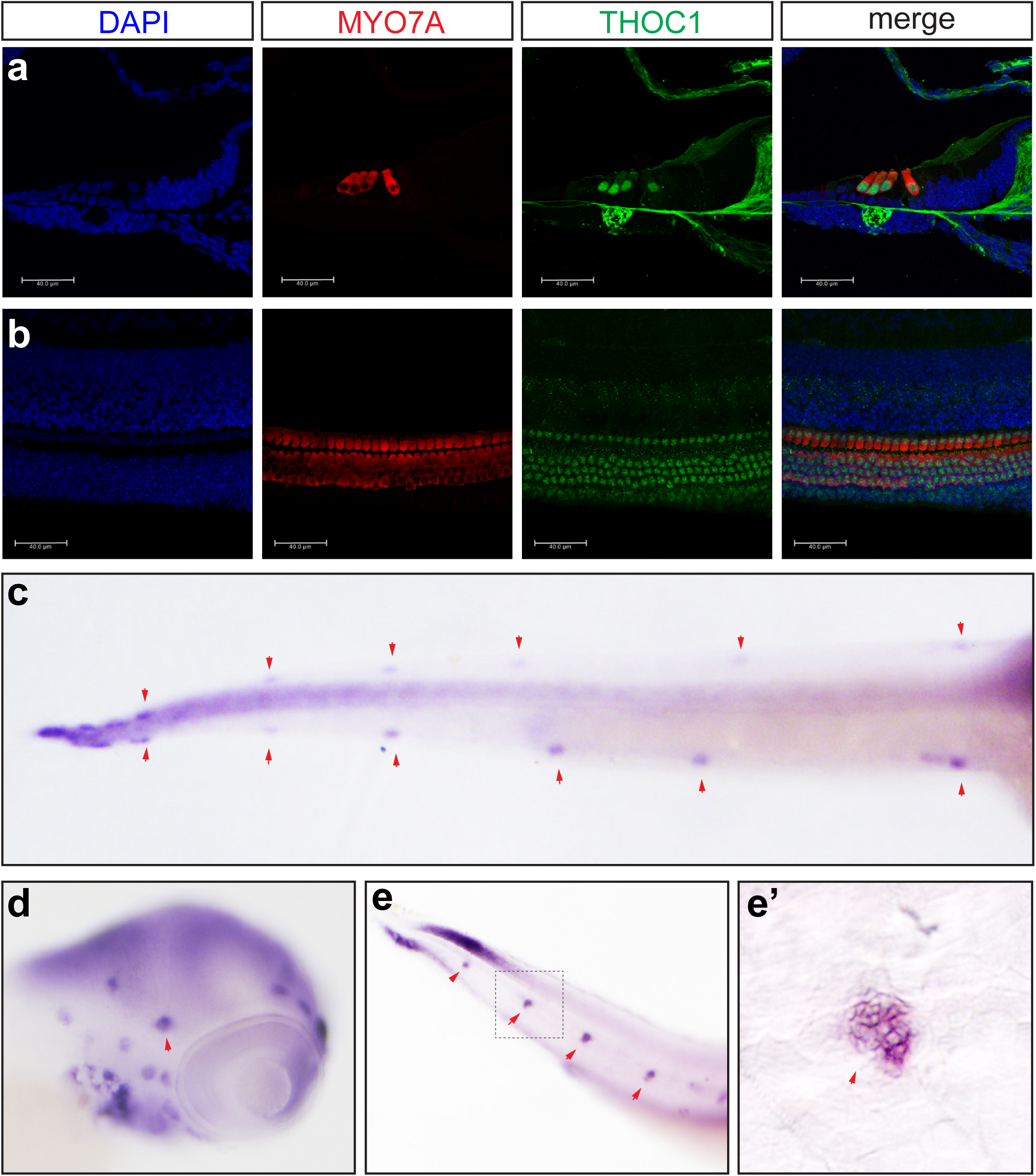
The expression of THOC1 in hair cells. (a, b) Confocal microscopic imaging analysis of THOC1 antibody staining in mouse cochlea hair cells. THOC1 was enriched in outer hair cells (OHC) and inner hair cells (IHC) in the P0 mouse cochlea. Blue: DAPI staining of the cell nuclei. Red: Myosin 7a staining marking hair cells. Green: THOC1. Bars, 40 μm. (c-e’) whole mount *in situ* hybridization analysis of expression of *thoc1* in 3 dpf zebrafish. (c) Dorsal view, arrowheads indicate neuromasts. (d) Lateral view, arrowheads indicate neuromasts. (e) Lateral view, arrowheads indicate neuromasts. (e’) Lateral view, arrowheads indicate neuromasts. The magnified region of square in (e).

### *Thoc1* deficiency caused hair cell developmental defects in zebrafish

In order to examine whether *thoc1* was required for the formation of hair cell, the CRISPR/Cas9 system was utilized to generate a series of *thoc1* mutants in *Tg(pou4f3:gap43-GFP)* transgenic zebrafish line, in which the membrane of hair cell is labeled with GFP[11]. To obtain viable but hypomorphic alleles, we chose the target sites at the exon 20, the second last exon of zebrafish *thoc1* (Supplementary Figure S4a). The selected gRNA-Cas9 system efficiently induced a series of frameshifting indels in the targeting sites that were predicted to truncating a small portion of the C-terminal protein (Supplementary Figure S4b). In the *thoc1* mutants, the number of hair cells clusters in neuromasts and the number of neuromasts are dramatically decreased (Figure 3a-d; Supplementary Figure S5). Meanwhile, the hair cell number in both each neuromast and in the whole embryo is significantly reduced as well (Figure 3e-j). In addition, we examined the fast escape reflex, the C-shaped startle response. It was found that the probability of the C-startle response in *thoc1* deficient zebrafish was significantly lower than that in control zebrafish embryo and adults, suggesting *thoc1* mutants have hearing problems (Supplementary Figure S6, S7). To verify these phenotypes in *thoc1* mutants were indeed caused by loss of function of *thoc1*, we examined whether *thoc1* knockdown could phenocopy the mutants. A splicing-blocking mophorlino (MO) was validated to efficiently interfere the *thoc1* pre-mRNA splicing, caused the exon3 (61 bp) deletion and lead to the reading frame shift (Figure 4a, b). Sanger sequencing analysis confirmed this result (data not shown). It was demonstrated that *thoc1* morphants had the same phenotypes with those in *thoc1* mutants (Figure 4c). In addition, both human *THOC1* mRNA and zebrafish *thoc1* mRNA significantly rescued the hair cell defects in *thoc1* morphants (Figure 4c, d). These results substantiate that hair cell defects were specifically caused by inactivation of *thoc1*.

**Figure 3.**
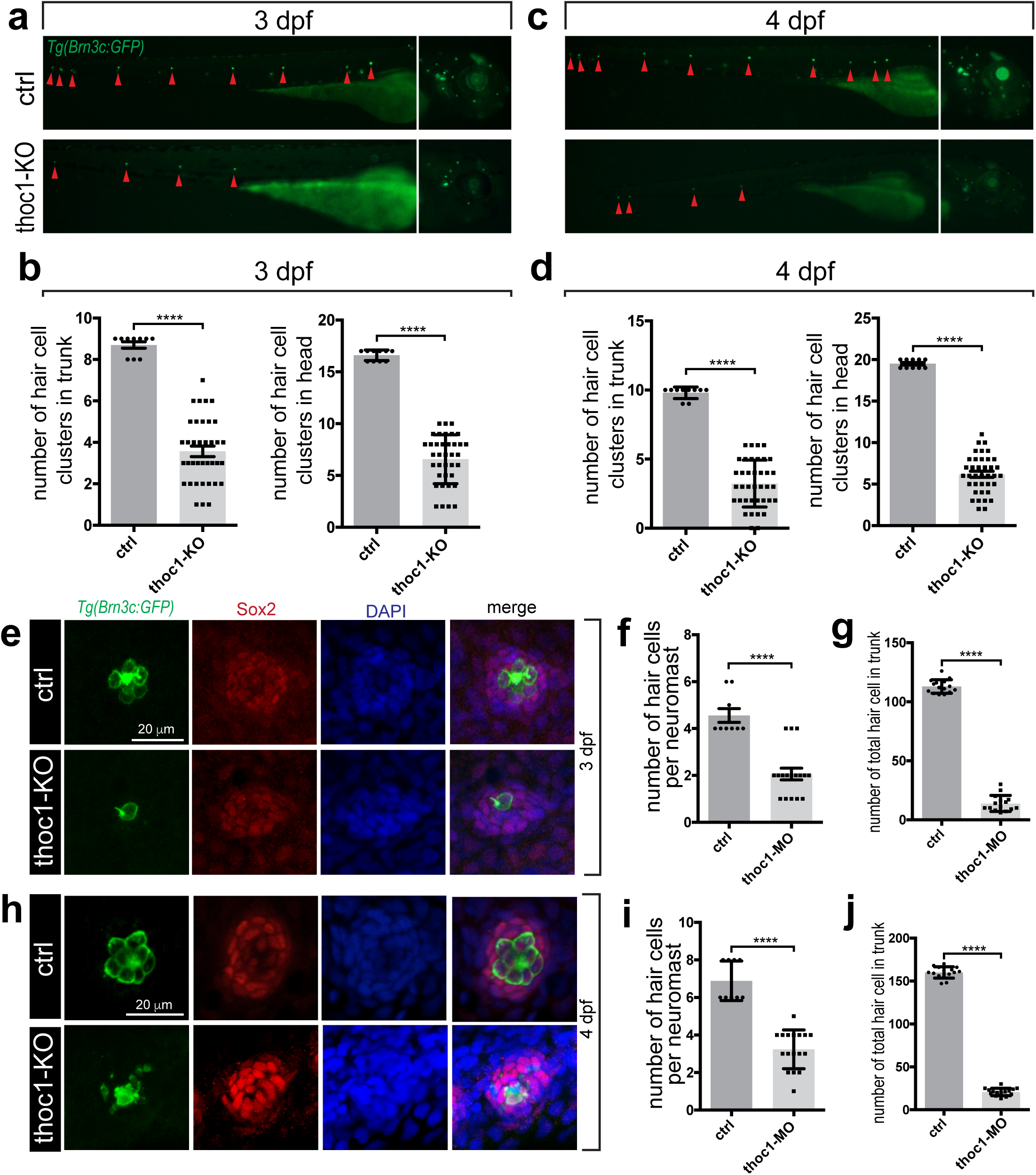
*Thoc1* deficiency caused hair cell developmental defects in zebrafish. (a) Fluorescence microscopic imaging analysis of *thoc1* mutant *Tg(pou4f3:gap43-GFP)* line at 3 dpf. Arrowheads indicate hair cell clusters. (b) Statistical analysis of the hair cell clusters in control and *thoc1* mutants (control, n=10; *thoc1* mutants, n=37). *t*-test, ****, *p*<0.0001. (c) Fluorescence microscopic imaging analysis of *thoc1* mutant *Tg(pou4f3:gap43-GFP)* line at 4 dpf. Arrowheads indicate hair cell clusters. (d) Statistical analysis of the hair cell clusters in control and *thoc1* mutants (control, n=10; *thoc1* mutants, n=38). *t*-test, ****, *p*<0.0001. (e) Confocal microscopic imaging analysis of the neuromasts in control and *thoc1* mutants at 3 dpf. (f, g) Statistical analysis of the hair cell number per neuromasts and total number in trunk of control and *thoc1* mutants (control, n=9; *thoc1* mutants, n=17, control, n=15; *thoc1* mutants, n=15) at 3 dpf. *t*-test, ****, *p*<0.0001. (h) Confocal microscopic imaging analysis of the neuromasts in control and *thoc1* mutants at 4 dpf. (i, j) Statistical analysis of the hair cell number per neuromasts and total number in trunk of control and *thoc1* mutants (control, n=9, *thoc1* mutants, n=17; control, n=15; *thoc1* mutants, n=15) at 4 dpf. *t*-test, ****, *p*<0.0001.

**Figure 4.**
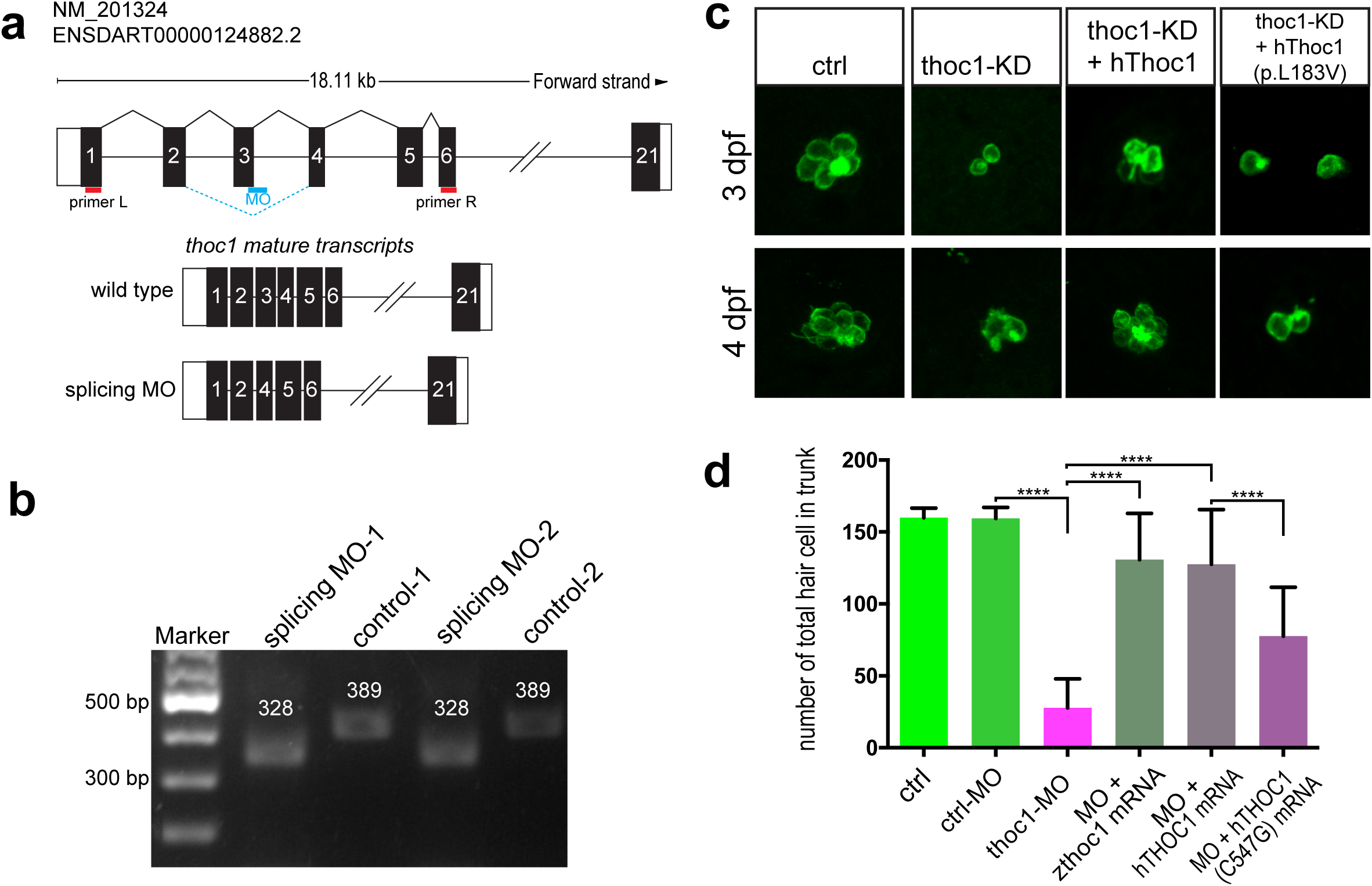
The THOC1 mutation (p. L183V) impaired its function in hair cell formation. (a) The diagram shows the targeting site of *thoc1* splice blocking morpholino and the RT-PCR primers design for validating the knockdown results. The wild type mature transcripts indicate the natural splicing product of *thoc1* mRNA. The splicing MO mature transcripts indicate the abnormal splicing product of *thoc1*mRNA with Exon3 deletion caused by morpholino injection. (b) The agarose gel electrophoresis image shows the 61bp Exon3 deletion. (c) Confocal microscopic imaging analysis of the hair cells in control, *thoc1* knockdown (KD), *thoc1* knockdown (KD) + *hThoc1* mRNA, and *thoc1* KD + *hThoc1* (p. L183V) mRNA at 3 dpf and 4 dpf. (d) Statistical analysis of the total hair cell number in trunk of control (n=15), control MO (n=15), *thoc1* MO (n=15), *thoc1* MO + *zthoc1* mRNA (n=15), *thoc1* MO + *hThoc1* mRNA (n=15), and *thoc1* MO + *hThoc1* (p. L183V) mRNA at 4 dpf (n=15). One-way ANOVA, ****, *p*<0.0001.

### The p. L183V mutation in *THOC1* impaired its function in hair cell formation

To testify whether the c.C547G/p.Leu183Val mutation in THOC1 impairs its function in hair cell formation, we explored the effect of this mutation during hair cell development. We setup 6 groups for microinjection and did confocal imaging analysis of hair cells at 4 days post fertilization (dpf): wild-type control, standard MO control, MO, MO + *zthoc1* mRNA, MO + *hTHOC1* mRNA, MO + *hTHOC1* (C547G) mRNA. The mutation of THOC1 significantly reduced the ratio of embryos been rescued by microinjection of *hTHOC1* (C547G) mRNAs into *thoc1* morphants (Figure 4c, d).

### Inactivation of *thoc1* induced apoptosis in neuromasts

To understand the cellular mechanism by which the hair cells were dramatically decreased, we did the confocal imaging analysis of the residual cells. In *thoc1* mutants, we found that around 83% of these hair cells had an abnormal morphology, such as distorted shape, quite smaller, and even became cell fractions (Figure 5a). These results suggest a number of hair cells underwent apoptosis. To test this hypothesis, we carried out Terminal deoxynucleotidyl transferase dUTP nick end labeling (TUNEL) analysis in *thoc1* mutants. It was revealed that *thoc1* deficiency resulted in dramatic hair cell and supporting cell apoptosis in neuromasts (Figure 5b).

**Figure 5.**
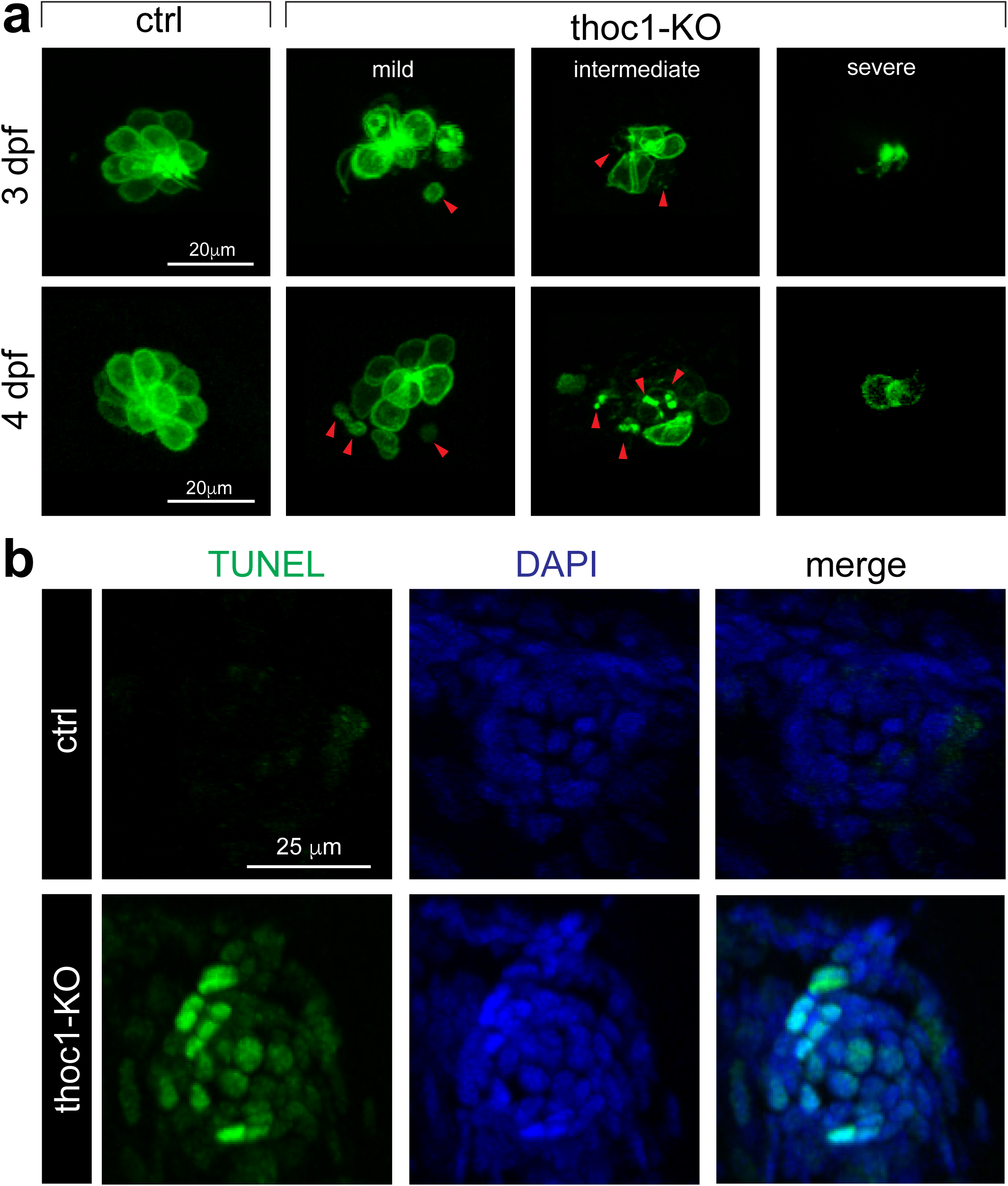
Inactivation of *thoc1* induced apoptosis in neuromasts. (a) Confocal microscopic imaging analysis of the hair cells in control and *thoc1* KO *Tg(pou4f3:gap43-GFP)* at 3 and 4 dpf, categorized into mild, intermediate, and severe groups. Arrowheads indicate the abnormal hair cells. (b) TUNEL analysis of the neuromasts in control and *thoc1* KO embryos.

In addition, we performed the transcriptome sequencing analysis of the *thoc1* mutated embryos at 48 hpf. We found that 455 genes were significantly up-regulated, while 1312 genes were significantly down-regulated in *thoc1* mutants compared to the sibling control (Figure 6a, b). KEGG analysis of these different expression genes revealed that 35 pathways were enriched, including P53 signaling pathway, DNA replication, and cell cycle (Figure 6c). The *p53* upregulation was validated by Realtime PCR analysis at 48 hpf and 72 hpf (Figure 6d). In addition, Realtime PCR analysis indicated that apoptosis genes *bax, casp3* and *casp9* were significantly up-regulated in 4 dpf *thoc1* mutants (Figure 6e). These results were in support of that inactivation of *thoc1* induced apoptosis in neuromasts.

**Figure 6.**
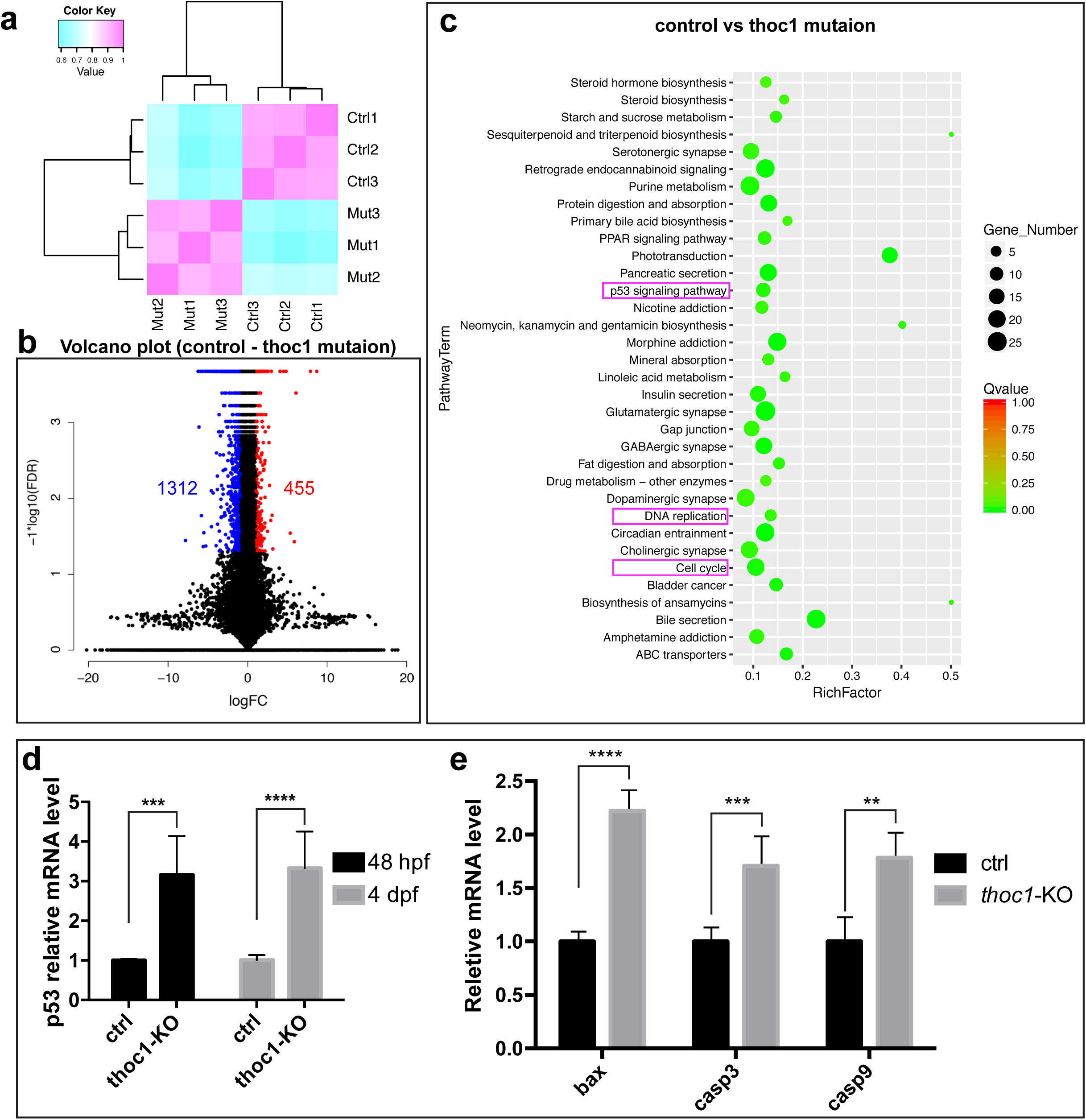
Transcriptome sequencing analysis of 48 hpf *thoc1* mutants. (a) Clustering analysis indicates the replicates within group have a good repeatability, while the control and mutated group are different. (b) Volcano diagram of different expression genes. Red dots indicate up-regulated genes; blue dots indicate down-regulated genes. Abscissa indicates gene fold change in different samples; ordinate represents statistical significance of gene expression change. (c) KEGG analysis plot of the differential gene, with the vertical axis representing the pathway and the horizontal axis representing the Rich factor. The size of the dot indicates the number of differentially expressed genes in the pathway, and the color of the dot corresponds to a different Qvalue range. (d) Relative mRNA levels of *p53* in control and *thoc1*-KO embryos at 48 hpf and 4 dpf (three times experiments, n=10 for each time). *t*-test; ***, *p*<0.001; ****, *p*<0.0001. (e) Relative mRNA levels of *bax, casp3* and *casp9* in control and *thoc1*-KO embryos at 4 dpf (three times experiments, n=10 for each time), *t*-test; **, *p*<0.01; ***, *p*<0.001; ****, *p*<0.0001.

To confirm the hair cell reduction in *thoc1* deficient embryos was due to the cell apoptosis, we knockdown the *thoc1* in *p53* mutated zebrafish embryos. It was revealed that depletion of *p53* partially restored the number of hair cells in *thoc1* morphants (Supplementary Figure S8). Additionally, inhibition of P53 by Pifithrin-α in the *thoc1* mutants significantly alleviated the *thoc1* induced apoptosis in neuromasts (Figure 7, Supplementary Figure S9).

**Figure 7.**
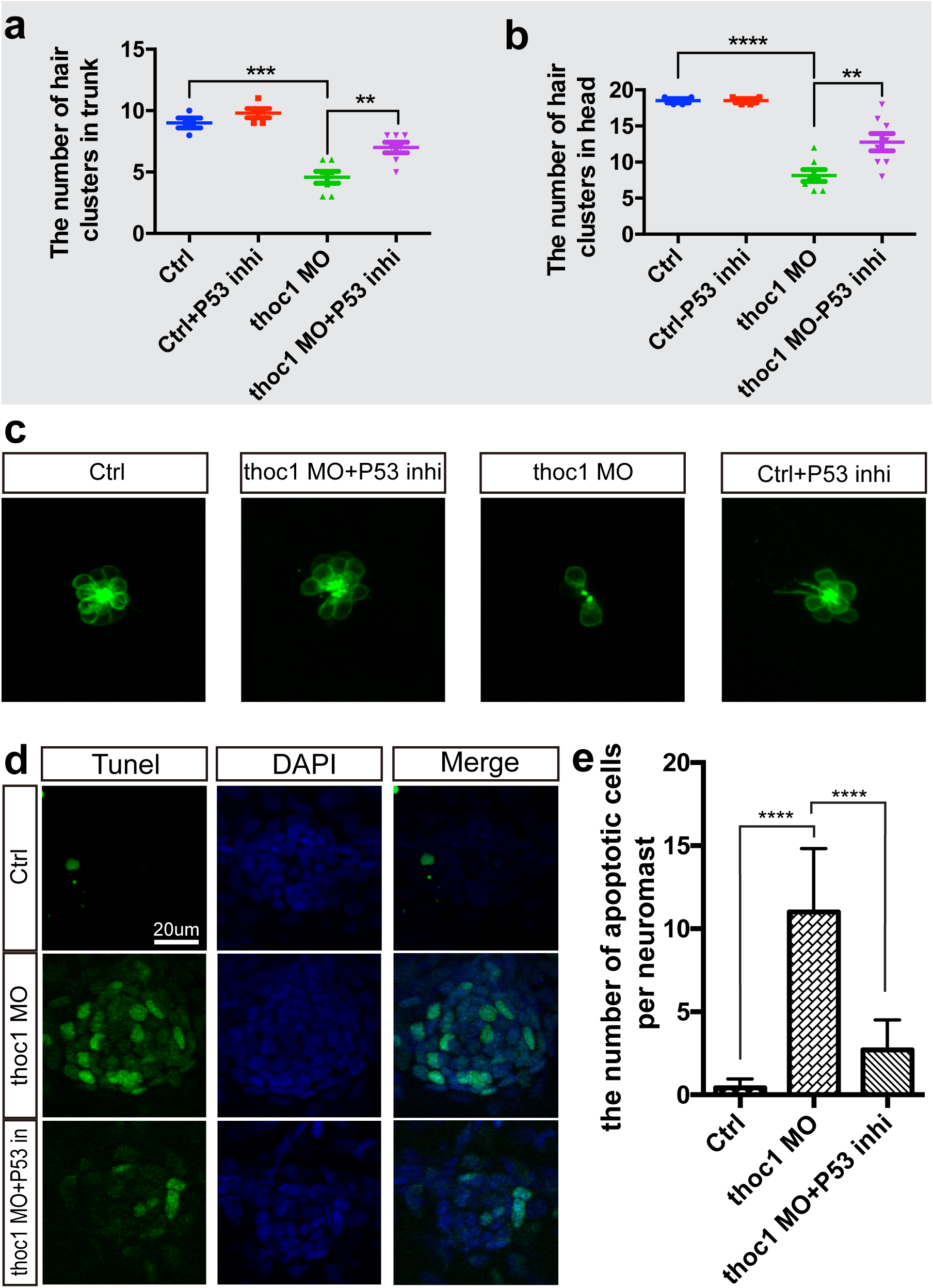
Inhibition of P53 signaling alleviated the *thoc1* induced apoptosis in neuromasts. (a, b) Statistical analysis of the hair cell clusters in control (n=8), ctrl + P53 inhibitor treatment (n=9), *thoc1* MO (n=8), and *thoc1* MO + P53 inhibitor treatment embryos (n=7). One-way ANOVA, ****, *p*<0.0001; ***, *p*<0.001; **, *p*<0.01. (c) Confocal microscopic imaging analysis of the hair cells in control, *thoc1* MO + P53 inhibitor treatment, thoc1 MO, and P53 inhibitor treatment *Tg(pou4f3:gap43-GFP)* embryos at 4 dpf. (d) TUNEL analysis of the neuromasts in control (n=7), *thoc1* MO (n=7), and *thoc1* MO + P53 inhibitor treatment embryos (n=7). (e) Statistical analysis of the number of apoptotic cells per neuromast in control, *thoc1* MO, and *thoc1* MO + P53 inhibitor treatment embryos. One-way ANOVA,****, *p*<0.0001.

## Discussion

As a key component of the TREX protein complex, THOC1 is evolutionarily conserved (supplementary Figure S2) and plays an essential role for coordinated gene expression during early and postnatal development [7, 9, 10]. In humans, non-synonymous variants in *THOC1* are extremely rare, as no such variants have minor allele frequencies higher than 0.002 in the Genome Aggregation Database (gnomAD, http://gnomad.broadinstitute.org/). In this study, we presented to our knowledge the first *THOC1* mutation associated with human mendelian disorders. Several lines of genetic and functional evidences supported that the p.L183V variant in *THOC1* is the probable cause of the late-onset, non-syndromic hearing loss in Family SH: a) *THOC1* was harbored in the 1.4-Mb critical interval defined by genome-wide linkage analysis with a very high maximum LOD score of 4.93 (Figure 1b and Supplementary Table S1, S2); b) Exome sequencing identified p.L183V in *THOC1* as the only candidate pathogenic variant segregating with the hearing phenotype in 21 members of Family SH (Figure 1a and Supplementary Table S2); c) the p.L183V variant changed a highly conserved amino acid (Figure 1e), was predicted to be pathogenic by computational programs Mutation Taster, PROVEAN and SIFT, and was not seen in 1000 ethnically matched normal hearing controls; d) the hair cell loss of the *thoc1*-knockdown zebrafish can be functionally rescued by embryonic microinjection of the wild-type *thoc1* mRNA, but to significantly lesser degree by that of the p.L183V mutant mRNA (Figure 4).

Even though THOC1 regulates expression of a wide range of genes required for crucial biological processes such as embryogenesis, organogenesis and cellular differentiation[8], mice with hypomorphic alleles in *Thoc1* were viable [10] and are associated with tissue-specific defects such as compromised fertility [9]. In our study, the affected members of Family SH showed no abnormalities other than the hearing loss, further demonstrating that systematic reduced level of functional Thoc1 is generally tolerated and may lead to certain tissue-restricted disorders.

To elucidate the role of THOC1 in the inner ear, we generated a series of *Thoc1* mutant zebrafish that mimicked the human hearing impairment by losing the C-startle response (Supplementary Figure S3, Figure 5, 6). These mutants likely represented hypomorphic alleles of *Thoc1* as the MO-knockdown zebrafish had similar phenotype (Figure 4). In inner ear, the most striking finding from both *Thoc1* mutant and knockdown zebrafish is the marked loss of hair cells due to apoptosis (Figure 3, 4, 5). Surprisingly, though hair cell apoptosis is a key step towards progressive and age-related hearing loss, *THOC1* represents only one of a very few deafness-causing genes directly involved in this process. Given THOC1 regulates distinct and specific sets of downstream genes in different tissues and development stages, the *Thoc1* deficiency zebrafish generated in our study presented a good model to study the coordinated control of hair cell apoptosis at the molecular level. Indeed, by transcriptome sequencing, we identified a list of genes that were differentially expressed in the *Thoc1* mutant zebrafish (Figure 6). Among them, genes in pathways associated with apoptosis such as p53 signaling, DNA replication and cell cycle were enriched.

Previous studies have suggested dual roles of Thoc1 in relation to cell proliferation and apoptosis depending on the context of different tissues and cell types. In embryonic development of *Rb1* null mice, Thoc1 is required to for increasing expression of *E2f* and other apoptotic regulatory genes[12]. On the other hand, Thoc1 is overexpressed in a variety of cancers[13, 14], and depletion of Thoc1 in those cancer cells, but not in normal cells, induce apoptotic cell death[15]. In mouse models, hypomorphic *Thoc1* allele lead to germ cell apoptosis that in testes correlates with the decreased number of primary spermatocytes[9]. In consistence with the latter findings, our study demonstrated that Thoc1 deficiency may induce hair cell apoptosis in zebrafish, which likely underlies the pathogenic mechanism of the late-onset, progressive hearing loss in Family SH. Accompanied with the Thoc1 deficiency was the increased expression of a number of downstream apoptotic genes such as *P53, Bax, Casp3* and *Casp9* (Figure 6). The p53 signaling pathway has been implicated in the hair cell apoptosis during age-related hearing loss[16]. In our study, we showed that inhibition of P53 by Pifithrin-α significantly alleviated the hair cell apoptosis in the *Thoc1* MO-knockdown zebrafish, presenting a potential new strategy for preventing of the age-related hearing loss.

In conclusion, our study identified *THOC1* as a new causative gene for late-onset, progressive hearing loss in humans. Functional studies showed that Thoc1 deficiency unleashes expression of proapoptotic genes in the p53 signaling pathway and results in hair cell apoptosis in zebrafish. These findings may provide important insight into the molecular basis of age-related hearing loss.

## MATERIALS AND METHODS

### Subjects and clinical evaluation

Members of Family SH was recruited through the Affiliated Hospital of Nantong University in Nantong, Jiangsu Province, China. A total of 9 affected and 18 unaffected members participated in the present study (Figure 1a, asteriskes). All subjects received comprehensive auditory evaluation including pure tone audiometry (PTA), otoscopic examination and temporal bone high-resolution CT scanning. Family history and general physical examination were performed to exclude the possible syndromic hearing loss. Informed consent was obtained from all participating subjects and this study was approved by the ethics committee of Affiliated Hospital of Nantong University.

### Genome-wide linkage analysis

Genome-wide multipoint linkage analysis was performed using the HumanOmniZhongHua-8 BeadChip (Illumina, San Diego, USA) and based on genotypes of 6301 Tag SNPs with an averaged 0.5cM resolution. Logarithm of the odds (LOD) scores were calculated by the parametric linkage analysis package Merlin v. 1.1.23. The inheritance model was set to dominant with full penetrance. The disease allele frequency was set to 0.0001.

### Whole-exome sequencing and verification of the pathogenic variants

Whole-exome sequencing was performed in four affected (III-4, III-9, III-10 and III-18) and two unaffected (III-16 and IV-6) members (marked with triangles in Figure 1A). Exons and flanking intronic regions of 20794 genes (33.2 Mb, 97.2% of cCDS coding exons), microRNAs and other non-coding RNAs were captured by Illumina TruSeq Exome Enrichment Kit and sequenced on a HiSeq 2000 instrument (Illumina, San Diego, CA, USA). Reads were aligned to NCBI37/hg19 assembly using the BWA Multi-Vision software package. SNPs and indels were identified using the SOAPsnp software and the GATK Indel Genotyper, respectively. Candidate pathogenic variants were defined as nonsense, missence, splice-site and indel variants with allele frequencies of 0.001 or less in public variant databases dbSNP, 1000 Genomes and previous sequencing data of 1000 Chinese Han adult normal hearing controls (in-house whole-exome sequencing data using the same platform). Candidate pathogenic variants were further genotyped in all family members by Sanger sequencing. Possible pathogenic effects of the identified mutation were evaluated by computational tools including Mutation Taster (http://www.mutationtaster.org), PROVEAN and SIFT (with cut-off scores set at -1.3 and 0.05, respectively, http://sift.jcvi.org).

### Immunostaining of THOC1 in mouse inner ear

Immunofluorescence staining of THOC1 was performed in cross-section and whole mount samples of mouse inner ear as previously described [17]. All procedures of the present study had approval from the Animal Use and Care Committee of Affiliated Hospital of Nantong University. Antibodies used in this study included mouse anti-THOC1 (SAB2702154, Sigma-Aldrich, St. Louis, MO), rabbit anti-MYOSIN VIIA (25–6790, Proteus Biosciences, Ramona, CA), and DAPI (D3571, Thermo-fisher, Waltham, MA). Immunostaining presented in figures was representative of two individual experiments.

### Zebrafish Line and Startle Response Test

The zebrafish embryos and adults were maintained in zebrafish Center of Nantong University under standard conditions in accordance with our previous procedures [18, 19]. The transgenic zebrafish lines *Tg(pou4f3:gap43-GFP)* and *Tg(cldnb:lynGFP)* were used as previously described [20]. Sound-evoked C-shaped startle response was tested at larval and adult stage in a well-plate and a plastic tank and recorded with a high-speed camera.

### RNA Isolation, Reverse Transcription (RT) and Realtime PCR

Total RNA was extracted from zebrafish embryos by TRIzol reagent according to the manufacturer’s instructions (Thermo Fisher Scientific, USA). Genomic contaminations were removed by DNaseI, and then 2 µg total RNA was reversely transcribed using a reversed first strand cDNA synthesis kit (Fermentas, USA) and stored at −20°C. The sequences of PCR primers, used for validating the Splice-blocking Morpholino, were: 5’-TCCGTCTCACTTCGACTTCA-3’ and 5’-TCCCAGCAGAGTAAAATGTGT-3’. The Realtime PCR analysis was performed according to the procedure described in our previous work [18]. The primers were used for the *bax*: 5’-AGAGGGTGAAACAGACCAGC-3’ and 5’-GCTGAACAAGAAAGGGCACAG-3’; for *caspase3*: 5’-TGGCACTGACGTAGATGCAG-3’ and 5’-GAAAAACACCCCCTCATCGC-3’; for *caspase9*: 5’-ACTAAATGACCGCAAGGGCT-3’ and 5’-TTGCCTCAGTGCCATGTGAA-3’; for *p53*: 5’-GCAAAAACTTGCCCCGTTCA-3’ and 5’-GCTGATTGCCCTCCACTCTT-3’.

### Whole Mount *in situ* Hybridization

A 498 bp cDNA fragment of Thoc1 was amplified from the cDNA library that established from wild type (WT) AB embryos using the specific primers of *thoc1* F1 5’-TGGAATCTGAACCCCGACAA-3’ and *thoc1* R1 5’-TGTCCGACTCGATCACTCTG-3’. Digoxigenin-labeled sense and antisense probes were synthesized using linearized pGEM-Teasy vector subcloned with this *thoc1* fragment by *in vitro* transcription with DIG-RNA labeling Kit (Roche, Switzerland). Zebrafish embryos and larvae were collected and fixed with 4% paraformaldehyde (PFA) in phosphate-buffered saline (PBS) for one night. The fixed samples were then dehydrated through a series of increasing concentrations of methanol and stored at −20°C in 100% methanol eventually. Whole mount *in situ* hybridization was subsequently performed as described in the previous study [21, 22].

### sgRNA/ Cas9 mRNA Synthesis and Injection

*Cas9* mRNA was obtained by in vitro transcription with the linearized plasmid pXT7-Cas9 according to the procedure previously described. The sgRNAs were transcribed from the DNA templates that amplified by PCR with a pT7 plasmid as the template, a specific forward primer and a universal reverse primer. The gRNA sequence is listed in the following: 5’-TAATACGACTCACTATAGGTGATTAAAACCGGAGAGGGTTTTAGAGCTAG AAATAGC-3’. *Cas9* mRNAs were synthesized *in vitro* using the linearized constructs as templates with SP6/T7 mMESSAGE mMACHINE Kit (Thermo Fisher Scientific, USA), purified with RNeasy Mini Kit (Qiagen, Germany), and dissolved in RNase free Ultrapure water (Thermo Fisher Scientific, USA). The sgRNAs were synthesized by the MAXIscript T7 Kit (Thermo Fisher Scientific, USA), and were purified with RNeasy Mini Kit (Qiagen, Germany), and dissolved in RNase free Ultrapure water (Thermo Fisher Scientific, USA). Zebrafish lines were naturally mated to obtain embryos for microinjection. One to two-cell stage zebrafish embryos was injected with 2–3 nl of a solution containing 250 ng/µl *Cas9* mRNA and 15 ng/µl sgRNA. At 24 h post fertilization (hpf), zebrafish embryos were randomly sampled for genomic DNA extraction according to the previous methods to determine the indel mutations by sequencing.

### Morpholino and mRNAs Injections

*Thoc1* splice-blocking Morpholino (MOs; Gene Tools, USA) sequence was 5′-AGTAAGCTGTGGACTCACTATCTGC -3′. The sequence of a standard control MO oligo was 5′-CCTCTTACCTCAGTTACAATTTATA-3′. The MOs were diluted to 0.3 mM with RNase-free water and injected into the yolk of one to two-cell stage embryos and then raised in E3 medium at 28.5°C. The cDNAs containing the open reading frame of the target genes were cloned into PCS2+ vector respectively and then was transcribed *in vitro* using the mMESSAGE mMACHIN Kit (Thermo Fisher Scientific, USA) after the recombinant plasmids linearized with NotI Restriction Enzyme (NEB, USA), and then the capped mRNAs were purified by RNeasy Mini Kit (Qiagen, Germany). 2 nl target genes mRNA were injected at 50 ng/µl into 1/2-cell stage embryos.

### FM1-43FX Labeling

To visualize and image the hair cells in lateral line neuromasts, the vital dye FM1-43FX (Thermo Fisher Scientific, USA) was applied at a concentration of 3 μM to live larvae for 45 s in the dark. After quickly rinsing three times with fresh water, the larvae were anesthetized in 0.02% MS-222 and fixed with 4% PFA in PBS for 2 h at room temperature or overnight at 4°C.

### TUNEL Staining

For TUNEL (Terminal deoxynucleotidyl transferase-mediated dUTP nick end labeling) assays, larvae were incubated in 0.1 M glycine/PBS solution for 10 min and then rinsed with PBT-2 three times for 10 minutes each. The larvae were then processed using the *In Situ* Cell Death Detection Kit (Roche, Switzerland) following the directions supplied by the manufacturer.

### mRNA sequencing by Illumina HiSeq

Total RNA of each sample was extracted using TRIzol Reagent (Invitrogen). RNA samples were quantified and qualified by Agilent 2100 Bioanalyzer (Agilent Technologies, USA). Next generation sequencing library preparations were constructed according to the manufacturer’s protocol (Illumina, USA). The libraries were sequenced using Illumina HiSeq instrument according to manufacturer’s instructions (Illumina, USA). The sequences were processed and analyzed by GENEWIZ. Differential expression analysis was carried out using the DESeq Bioconductor package. After adjusted by Benjamini and Hochberg’s approach for controlling the false discovery rate, P-value of genes were set <0.05 to detect differential expressed ones.

### Microscopy and Statistical Analysis

Zebrafish embryos were anesthetized with E3/0.16 mg/mL tricaine/1% 1-phenyl-2-thiourea (Sigma, USA) and embedded in 0.8% low melt agarose, and then were examined with a Leica TCS-SP5 LSM confocal imaging system. For the results of *in situ* hybridization, Photographs were taken using an Olympus stereomicroscope MVX10. All images of THOC1 immunostaining in mouse were imaged using the Leica SP5 confocal microscope. Statistical analysis was performed using GraphPad Prism software. T-test and one-way analysis of variance (ANOVA) were used, and P < 0.05 were considered statistically significant.

## Author contributions

DL, TY, LPZ, and HW conceived the project. LPZ, TY and FFS did the linkage and sequencing analysis. LPZ and FFS contributed to the collection of clinical specimens and the analysis of human data. YG, RZ, FFS, CC, FPQ, XCD, GYW and RJC carried out the zebrafish and mouse experiments. LPZ, DL and YG contributed to the design and interpretation of the experiments. PHC and XHP carried out the normal hearing screening experiment. LPZ, TY and DL wrote the manuscript and all authors commented and approved the manuscript.

## Acknowledgements

This work was supported by National Natural Science Foundation of China (81641155 and 81870725 to LP.Z; 81371101 and 81570930 to TY; 81330023 to HW; 81570447 and 81870359 to DL; 81700920 to XP and 81600802 to RZ), Shanghai Municipal Science and Technology Commission (14DZ2260300 to HW) and Shanghai Municipal Education Commission-Gaofeng Clinical Medicine Grant (20152519 to TY), Nantong science and technology plan frontier and key technology innovation fund (MS22015048 to LP.Z); and the Foundations from Jiangsu Province (BK20180048 and 17KJA180008) to DL.

